# Existing host range mutations constrain further emergence of RNA viruses

**DOI:** 10.1101/394080

**Authors:** Lele Zhao, Mansha Seth Pasricha, Dragos Stemate, Alvin Crespo-Bellido, Jacqueline Gagnon, Siobain Duffy

## Abstract

RNA viruses are capable of rapid host shifting, typically due to a point mutation that confers expanded host range. As additional point mutations are necessary for further expansions, epistasis among host range mutations can potentially affect the mutational neighborhood and frequency of niche expansion. We mapped the mutational neighborhood of host range expansion using three genotypes of the dsRNA bacteriophage phi6 (wildtype and two isogenic host range mutants) on the novel host *Pseudomonas syringae* pv. *atrofaciens* (PA). Sanger sequencing of fifty PA mutant clones for each genotype and population Illumina sequencing both revealed the same high frequency mutations allowing infection of PA. Wildtype phi6 had at least nine different ways of mutating to enter the novel host, eight of which are in p3 (host attachment protein gene), and 13/50 clones had unchanged p3 genes. However, the two isogenic mutants had dramatically restricted neighborhoods: only one or two mutations, all in p3. Deep sequencing revealed that wildtype clones without mutations in p3 likely had changes in p12 (morphogenic protein), a region that was not polymorphic for the two isogenic host range mutants. Sanger sequencing confirmed that 10/13 of the wildtype phi6 clones had nonsynonymous mutations in p12 and two others had point mutations in p9 and p5 – none of these genes had previously been associated with host range expansion in phi6. We demonstrate, for the first time, epistatic constraint in an RNA virus due to host range mutations themselves, which has implications for models of serial host range expansion.

**Importance:** RNA viruses mutate rapidly and frequently expand their host ranges to infect novel hosts, leading to serial host shifts. Using an RNA bacteriophage model system (*Pseudomonas* phage phi6), we studied the impact of pre-existing host range mutations on another host range expansion. Results from both clonal Sanger and Illumina sequencing show extant host range mutations dramatically narrow the neighborhood of potential host range mutations compared to wildtype phi6. This research suggests that serial host shifting viruses may follow a small number of molecular paths to enter additional novel hosts. We also identified new genes involved in phi6 host range expansion, expanding our knowledge of this important model system in experimental evolution.

## Introduction

Emerging and re-emerging viruses that host shift to infect new species pose significant economic and health costs to humans, animals, plants and our ecosystems (1-3). While ecological exposure is an essential part of emergence on a novel host (2), spillover infection of the novel host typically requires a host range mutation – the genetic component of host range expansion (4). These exaptive host range mutations must exist in the viral population prior to contact with the novel host, as part of the virus’ standing genetic diversity (5, 6). The exact mutations and mechanisms of host shifting are intensively studied in emerging zoonotic viruses such as influenza, SARS-CoV, and Ebola virus (7, 8).

Given the high mutation rates (9), potentially large population sizes and fast replication of many emergent RNA viruses (10), they are capable of generating and maintaining substantial genetic variation (11, 12). This variation fuels adaptation, and selective sweeps leave genetic marks of past ecological history in viral genomes. These fixed mutations can alter the fitness landscape and constrain evolutionary trajectories of viruses due to epistatic interactions between mutations (13). Virus evolution is known to be shaped by epistasis, detected by both laboratory experimentation and phylogenetic analysis (14-16), and increased understanding of epistasis promises to improve our predictions of why some viral emergence events are more successful than others (17).

Some emergent viruses experience several hosts, often due to serial emergence events (18). MERS-CoV is proposed to have host jumped from its natural reservoir (bats) into camels, then later spilling over to the human population (19). Similarly, canine parvovirus jumped from infecting cats to raccoons and then jumped again to infect dogs (20). Influenza strains have also serially shifted hosts (*e.g*., H3N8 originated from avian hosts infecting horses, and then shifting to dogs (21)). This kind of serial emergence allows for the possibility of host range mutations themselves to play a significant role in shaping the landscape of further emergence – one of the legacies of previous host use (7, 22). We used the model RNA virus, *Pseudomonas* dsRNA bacteriophage phi6, to investigate the role of extant host range mutations on further host range expansion.

Phi6 has been a popular model for understanding host range mutations and their fitness effects (5, 23, 24). However, all previous studies have exclusively looked at a wildtype genotype, replicating in its reservoir host, instead of investigating the interactions of multiple host range mutations during frequent host shifting, or serial emergence. In this study, we mapped the host range mutational neighborhoods of wildtype phi6 and two isogenic host range mutants (E8G in P3; G515S in P3) emerging in novel host *P. syringae* pv. *atrofaciens* (PA). Significant epistatic constraint was observed with both host range mutants in clonal and Illumina sequencing – only one or two mutations were found that allowed infection of PA. These mutations were a subset of the large mutational neighborhood of PA host range mutations available to wildtype phi6. Additionally, we have identified host range associated genes other than the canonical site of host range mutations, three genes on the Small segment, not previously implicated in host-shifting. Our work supports using deep sequencing to map mutational neighborhoods in future studies, though both deep sequencing and the more labor-intensive characterization of clones complemented each other. This work provides a panoramic view of host range mutational neighborhoods in phi6, while demonstrating a significant constraint imposed by host range mutation in a fast-evolving RNA virus.

## Material and Methods

### Strains and culture conditions

Wildtype phi6 (ATCC no. 21781-B1) and its standard laboratory host: *P. syringae* pathovar *phaseolicola* (PP) strain HB10Y (ATCC no. 21781) were originally obtained from American Type Culture Collection (ATCC, Bethesda, MD). These, along with novel hosts *P. syringae* pathovar *atrofaciens* (PA), *P. syringae* pathovar *tomato* (PT), *P. pseudoalcaligenes* East River isolate A (PE) were streaked from glycerol stocks originally obtained from G. Martin (Cornell University, Ithaca, NY), and L. Mindich (Public Health Research Institute, Newark, NJ) as described in previous studies (23, 25). Previously isolated isogenic host range mutants phi6-E8G and phi6-G515S (with E8G and G515S mutation on the host attachment protein P3, respectively) (23) were used to examine genome-wide mutations on host expansion. Both host range mutants can infect PP, PT and PE. Bacteria were grown in LC media (LB broth pH 7.5), 25°C. Phages were grown with bacteria in 3mL 0.7% agar top layer on 1.5% agar plates as previously described (23).

### Mutational neighborhood mapping

Twice-plaque-purified phi6-WT, phi6-E8G, and phi6-G515S were raised to high-titer lysates on their respective hosts (i.e. phi6-WT was grown on PP while phi6-E8G and phi6-G515S on PT), and titered on their respective hosts. All lysates were tested for existent PA host range by spot plating approximately 10^4^-10^5^ plaque forming units (pfu) on a lawn of PA before plating to select for host range mutants. At least 10^6^ pfu of phage were plated on novel host PA to isolate one host range mutant per lysate. 50 single plaques were isolated from each lysate by plating on PA. Five of the phi6-G515S PA host range mutant plaques were isolated by the Biotechnology class of Spring 2016 at South Brunswick High School. All 150 plaques were stored in 40% glycerol at −20°C as freezer stock and generated into high-titer lysates again for further analyses.

### PA mutation frequency assays

Four independent clones of twice-plaque purified phi6-WT, phi6-E8G, and phi6-G515S plaques were raised to high-titer lysates on host PP, PT and PT respectively. After measuring titers on these hosts, these high titer lysates were titered on PA to assess the PA mutation frequency within the population standing genetic diversity. One of the four clones tested for each genotype was the source of the 50 clones.

### Fitness assays

Equal amounts of host range mutant were mixed with a common competitor (phi6-WT) in paired growth assays (PGA) (26) to test the mutant’s relative fitness on PP. Ratios of host range mutant and common competitor (CC) in the mixtures were obtained by counting the pfu of the initial mix (Day0) and after 24 hours of growth (Day1). The relative fitness to the common competitor was calculated using the following formula.

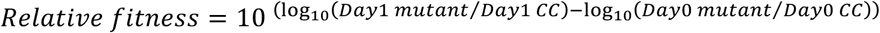

To distinguish the different genotypes, two hosts were mixed (20:1 PP:PA) to generate the bacterial lawn as phi6-WT can only infect PP, which creates turbid plaques while the mutants can infect both hosts creating clear plaques. Statistical analyses of fitness data, including ANOVA and Tukey’s honestly significant difference (HSD) tests, were performed in R (27).

### Sequencing

One microliter of the host range mutant glycerol stock was plated on a lawn of PA to generate high-titer lysates. Viral RNA was extracted from these lysates using QiaAmp Viral RNA Mini Kit (Qiagen, Valencia CA) per manufacturer guidelines. RT-PCR was conducted using SuperScript II Reverse Transcriptase (Invitrogen, now Thermo Fisher Scientific, MA) with random hexamers and KAPA Taq DNA Polymerase (Kapa Biosystems, now Roche) with primers that amplified the regions encoding P3 (host attachment) and P6 (membrane fusion) on the Medium segment. Amplified PCR products were cleaned up using EXO-SapIT (US Biological, Swampscott, MA) and Sanger sequencing was performed by Genewiz, Inc. (South Plainfield, NJ). Sequencing results were aligned and mutations identified with Sequencher 4.10.1.

### Library preparation

Phi6-WT, phi6-E8G and G515S were raised on their most recent host to obtain high-titer lysates, as described above. Each high-titer lysate was diluted and plated on PA to obtain a plate comprised of ˜400 plaques each. These plates were harvested to make lysates of phi6-WT, phi6-E8G and phi6-G515S capable of infecting PA. Viral RNA extracted using QiaAmp Viral RNA Mini Kit (Qiagen, Valencia CA) were purified by 1% low melt agarose gel electrophoresis (IBI Scientific, IA) and Gelase digestion following manufacturer instructions (GELase(tm) Agarose, Lucigen). Individual RNA samples at a final concentration of ˜15 ng/uL were prepared into Illumina RNA libraries using TruSeq RNA Library Prep Kit (Illumina, CA). Single-ended 150-cycle deep sequencing was performed on Illumina MiSeq housed in Foran Hall, Rutgers University (SEBS Genome Cooperative).

### NGS data analysis

Raw reads were trimmed and filtered with cutadapt 1.12 (Q score cutoff: 30, minimum length cutoff: 75bp, adapters and terminal Ns of reads removed) (28). Then, the reads were mapped to the *Pseudomonas* bacteriophage phi6 genomes (reference sequences derived from the Illumina sequencing of phi6-WT, phi6-E8G and phi6-G515S (also confirmed with Sanger sequencing)), using BWA-MEM with default settings (29). Although approximately 33.75%-66.88% of the NGS reads mapped to *Pseudomonas* host genome, all virus genome positions had above 1000X reads coverage, with the exception of the Large segment of phi6-G515S, which had 322X to 12,058X coverage. Additional file conversion was performed using SAMtools (30). Genome nucleotide counts by position were counted with Integrative Genomics Viewer IGVTools (count options: window size 1 and --bases) (31). Whole genomes variant calling was performed using VarScan (32). Shannon Entropy was calculated for each position of the genome with the following equation:

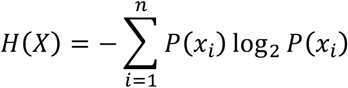

where n=4 for 4 nucleotides, *P*(*x*_*i*_) is the proportion of a single nucleotide over all nucleotides read at that position. When comparing levels of polymorphism between populations directly, the SNPs in each protein-coding gene were considered as independent observations for a paired t-test (Microsoft Excel, Redmond, WA).

## Results

### Mapping P3 PA mutational neighborhood

We found that PA host range mutational neighborhood is highly genotype-dependent. Fifty host range mutant plaques were isolated for each of the three genotypes (phi6-WT, phi6-E8G and phi6-G515S; Table 1) and their p3 genes for the phi6 attachment protein were Sanger sequenced. We only sequenced the p3 gene because P3 is the only highly accessible protein on the outside of the virion (33) and the only protein associated with phi6 host range in all previous studies (5, 23, 24). Thirty-five out of the fifty sequenced phi6-WT p3 sequences had single nonsynonymous mutations (seven unique mutations identified), two had double mutations, and thirteen had no detectable mutations on p3 gene. Forty-eight of the fifty phi6-E8G host range mutants contained one of the two single mutations present in the phi6-WT clones (A133V, S299W), the remaining two clones had double mutations: A133V and an additional nonsynonymous mutation. All 50 phi6-G515S host range mutants had the A133V mutation; 43 as a single mutation, five as double mutants and two as triple mutants. The nonsynonymous mutation A133V was the most frequent in the three tested populations: 24% in phi6-WT isolates, 96% in phi6-E8G isolates, and 100% in phi6-G515S isolates, making this the most prevalent mutation conferring infection of PA. In addition, there was a noticeable drop in diversity of single host range mutations from the phi6-WT population compared to the phi6-E8G and phi6-G515S populations, consistent with epistatic constraint on mutational neighborhood by host range mutations. We have summarized these P3 host range mutational neighborhoods on PA in a two-dimensional schematic (Figure 1).

**Table 1.**
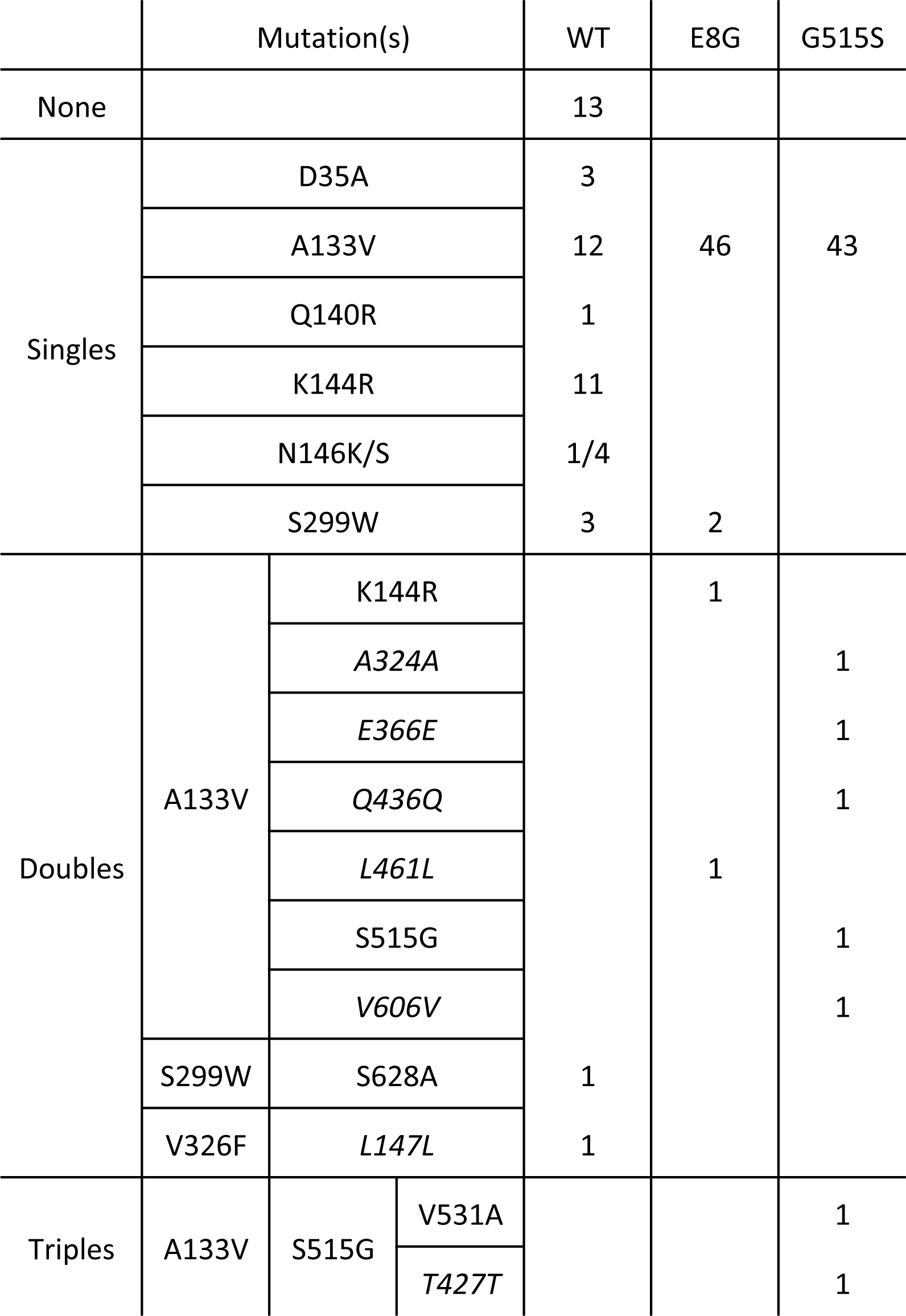
PA host range mutations detected in host attachment protein P3 with Sanger sequencing

**Figure 1.**
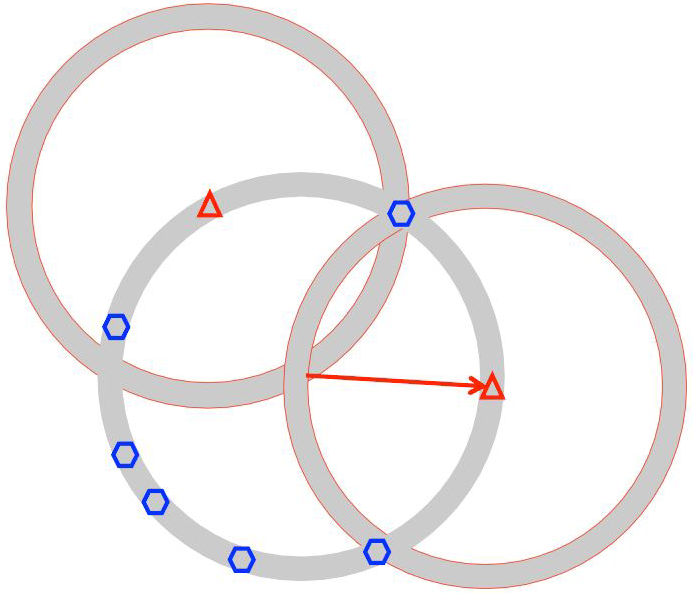
2D schematic representing mutational neighborhoods (shaded circles) of phi6 P3. Circles represent the P3 mutational neighborhood of the mutants, which are the centers of circles. The geometric shapes are known P3 mutants. The arrows are mutation events, such as host range expansion.

Several published works have investigated the mutational neighborhoods of phi6 p3 during expansion of host range (Table 2). Two sites were favored by host range expansion events onto *P. pseudoalcaligenes* and *P.syringae* pv *glyclinea*: the 8^th^ and 554^th^ amino acid of attachment protein P3 (66/81 and 18/39 of isolated mutants, respectively, (5, 24)). However, the PA mutational neighborhood does not include these frequent sites of mutation to other hosts. There is some overlap with previous studies, for instance N146S (5) and A133V (23), but this suggests that phi6 may interact differently with host PA during attachment than with other *Pseudomonas* species or *P. syringae* pathovars. The absence of host range mutations in p3 for thirteen of the phi6-WT PA isolates was unexpected, since 97% of previously independently isolated host range mutants had nonsynonymous mutations in p3, with no other sites in the phi6 genome identified as causing the expanded host range in the remaining 4/118 (5, 23, 24). This motivated a more in-depth approach: deep sequencing to map the entire mutational neighborhood.

**Table 2.** Host range mutational neighborhoods inferred from non-synonymous mutations of P3 from previous publications. Amino acids are in upper case letters with specific site in protein. Numbers indicate occurrence out of isolates sequenced. “Total” is the number of wild type isolates studied. *P. syringae* pv. *atrofaciens* and *P. syringae* pv. *glycinea* are closely related to the original host *P. syringae* pv. *phaseolicola*. While *P. pseudoalcaligenes* ERA is distantly related to the original host. (glycinea data from (5), ERA data from (24)).

| P3 aa mutation | *P. syringae* pv. *atrofaciens* | *P. syringae* pv. *glycinea* | *P. pseudoalcaligenes* ERA |
| --- | --- | --- | --- |
| G5S |  | 2 |  |
| E8K/G/D/A |  | 6 | 52 |
| D35A | 3 |  |  |
| Q130R |  |  | 1 |
| A133V | 12 |  |  |
| Q140R | 1 |  |  |
| K144R | 11 |  |  |
| D145G |  | 3 |  |
| N146K/S | 5 | 6 |  |
| E178D |  | 2 |  |
| S299W | 3 |  |  |
| V326F | 1 |  |  |
| P339H |  | 1 |  |
| T516A |  | 4 |  |
| D533A |  | 1 |  |
| D535N |  | 1 |  |
| D554G/A/V/N |  | 12 | 3 |
| L555F |  | 1 |  |
| Double/Triples | 1 |  | 10 |
| None | 13 | 1 | 3 |
| Total | 50 | 40 | 69 |

### Deep sequencing of phi6 populations

Each phi6 population was raised to high titer on its most recent host (phi6-WT on PP and phi6-E8G, phi6-G515S both on PT) and plated on PA to obtain a lysate made of ∼400 host range mutant plaques. All population lysates from before and after PA host range expansion were sequenced. Change in Shannon entropy was calculated to determine the sites that became more or less variable after overnight growth on PA. The signals of increased variation in the p3 gene matched Sanger sequencing results; all single mutations identified in the three genotypes underwent noticeable entropy change after gaining PA host range (Figure 2). We also found several sites in P3 that may have evaded detection by clonal sampling; including amino acid 247 of phi6-WT and amino acid 35 of phi6-G515S. Deep sequencing also revealed sites of high entropy change in other genes in phi6-WT that could be additional genes controlling host range, and suggesting targets for sequencing in the phi6-WT clones that did not contain p3 mutations (Figure 3, 4). Our results suggested that non-structural protein genes p12 (encoding the morphogenic protein) and p9 (encoding the major membrane protein) were the most probable sites of additional host range mutations; both genes are involved with viral nucleocapsid vesiculation of the host inner membrane (34, 35). Results from deep sequencing the p3 gene and from the entire genome further confirmed the constrained neighborhood of host range mutants phi6-E8G and phi6-G515S revealing fewer possibilities for PA mutations in p3, and none elsewhere in the genome.

**Figure 2.**
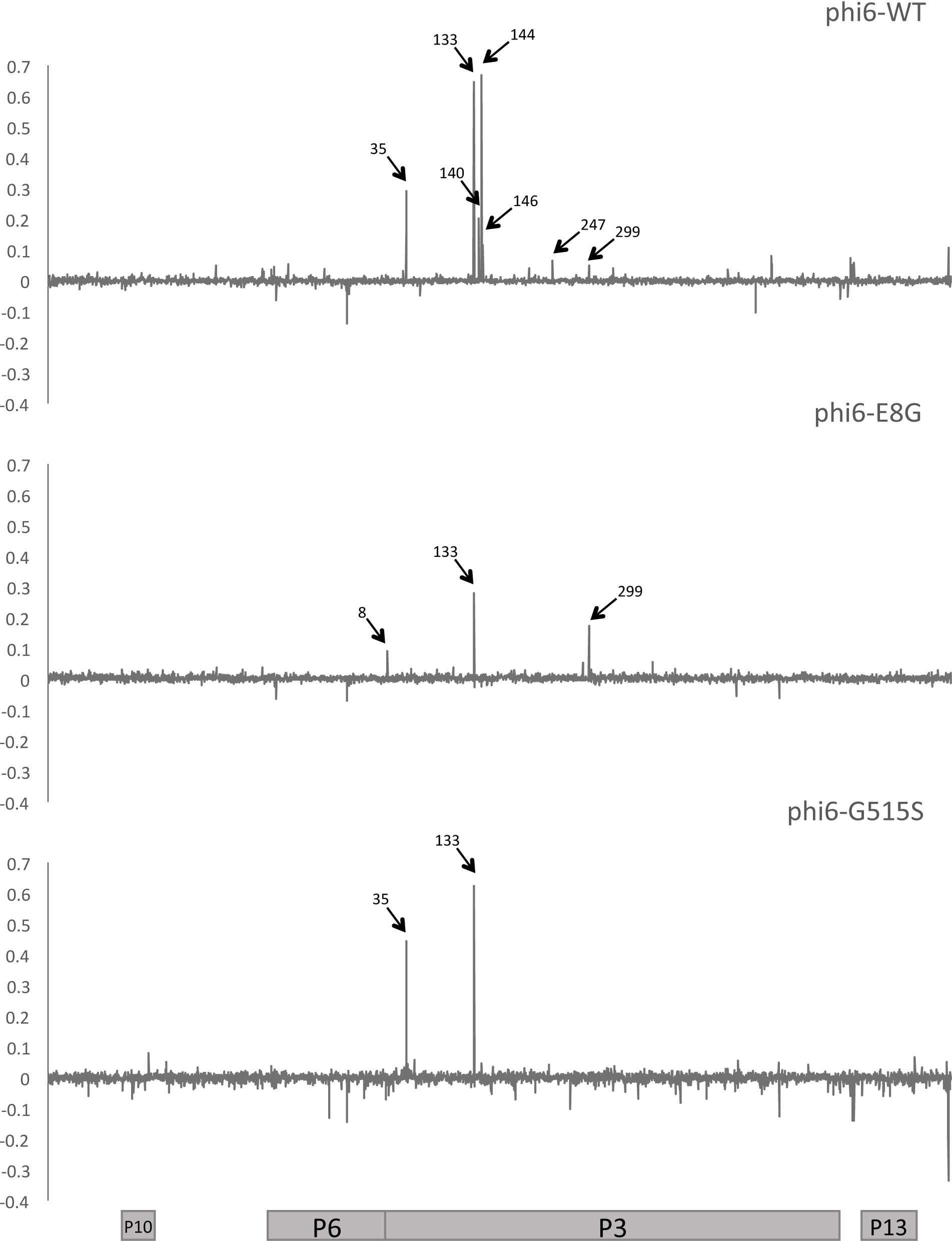
Change in Shannon Entropy in Medium segment of phi6-WT, phi6-E8G and phi6-G515S. Positions labeled correspond to amino acid position in P3. Coding regions of the Medium segment are aligned to the graphs.

**Figure 3.**
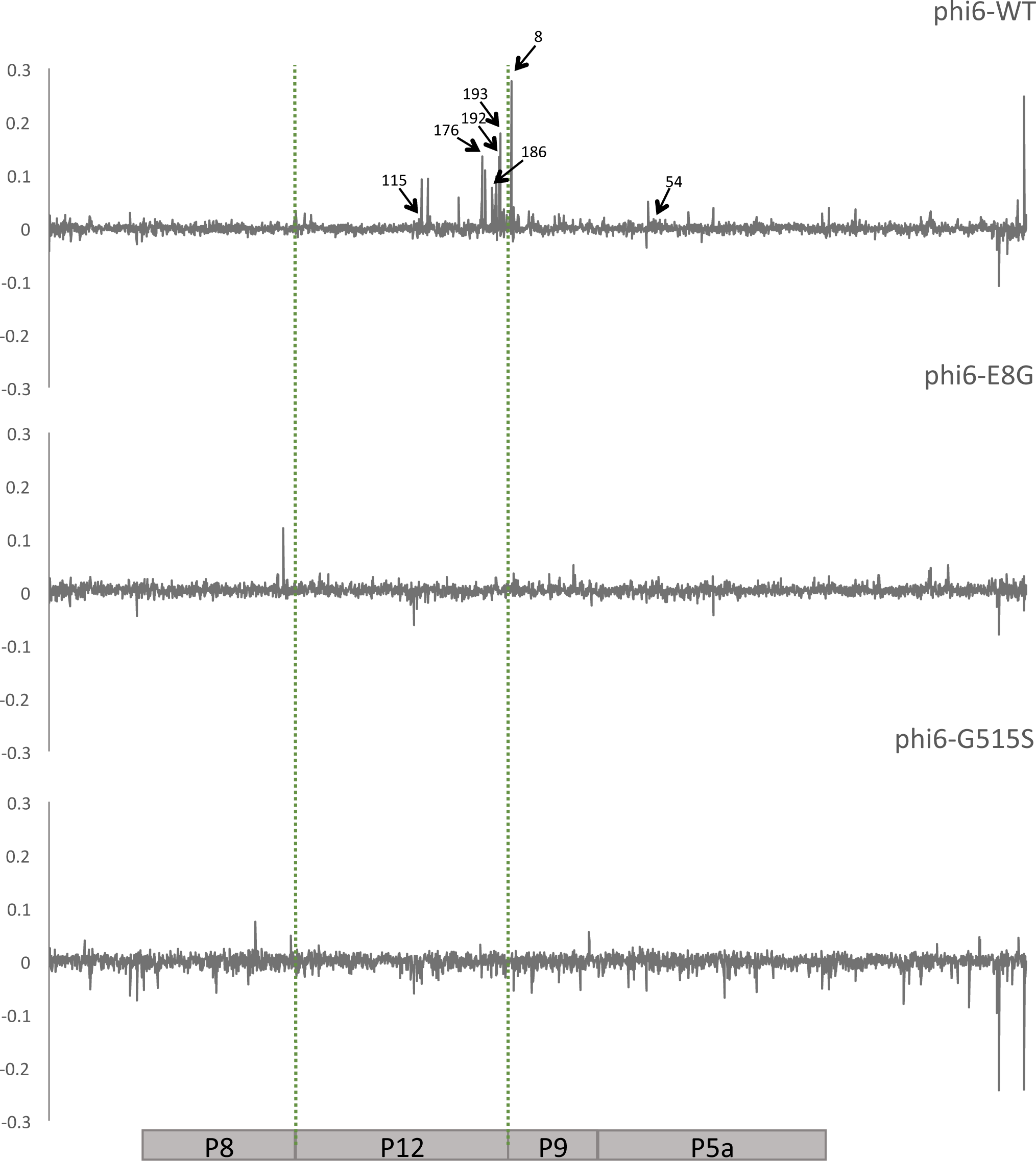
Change in Shannon Entropy in Small Segment of phi6-WT, phi6-E8G and phi6-G515S. Coding regions of the Small segment are aligned to the graphs. Positions labeled correspond to amino acid positions in aligned genes.

**Figure 4.**
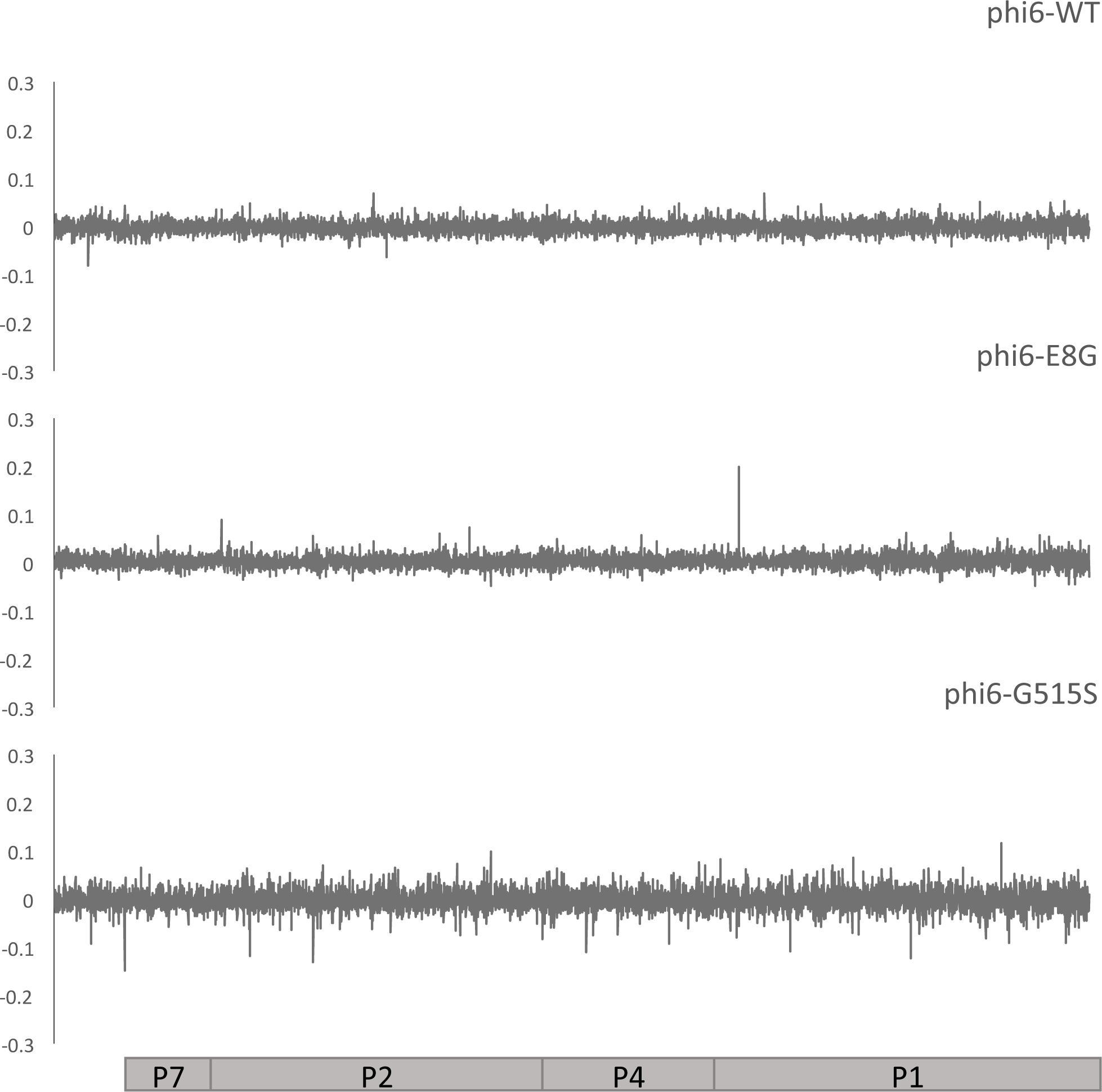
Change in Shannon Entropy in Large segment of phi6-WT, phi6-E8G and phi6-G515S. Coding regions of the Large segment are aligned to the graphs.

### Non-p3 phi6-WT mutant sequencing

We amplified and Sanger sequenced the Small segment of all phi6-WT isolates that did not show mutation in p3 (Table 3). Ten of the thirteen isolates contained a single nonsynonymous mutation in p12. One contained a single nonsynonymous mutation in p9, another contained a nonsynonymous mutation in p5. The final mutant had a single synonymous mutation in p9. This clone was then fully Sanger sequenced, but no nonsynonymous mutations were identified. These results matched many of the sites with the highest change in Shannon entropy we observed on the Small segment of deep sequenced phi6-WT populations, and strongly suggest that mutations in these non-structural genes can affect phi6 host range.

**Table 3.**
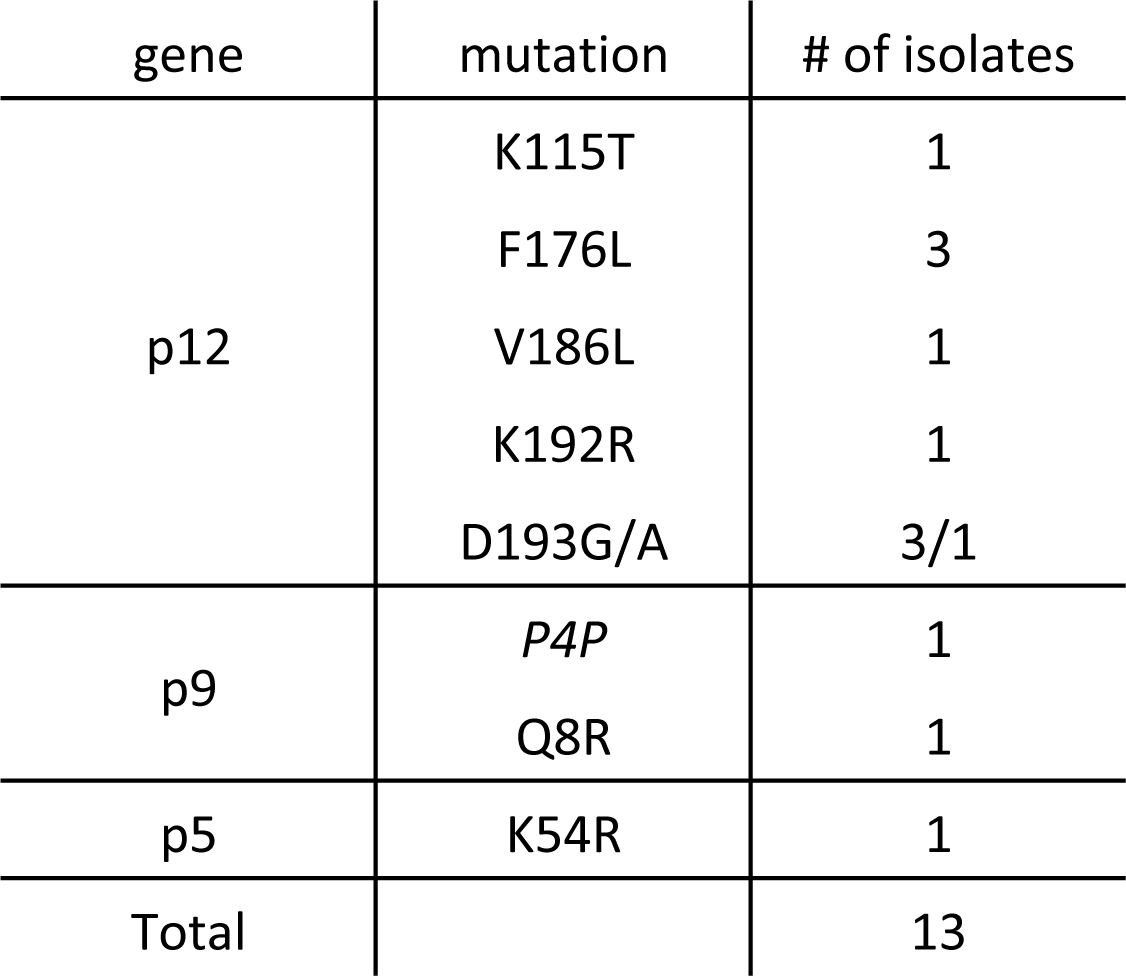
PA host range mutations detected on the small segment with Sanger sequencing.

| gene | mutation | # of isolates |
| --- | --- | --- |
| p12 | K115T | 1 |
|  | F176L | 3 |
|  | V186L | 1 |
|  | K192R | 1 |
|  | D193G/A | 3/1 |
| p9 | *P4P* | 1 |
|  | Q8R | 1 |
| p5 | K54R | 1 |
| Total |  | 13 |

### SNPs of host range expanded phi6 populations

In addition to the sites of highest entropy change, we called SNPs present in the deep sequenced pairs of populations. The counts and details of unique synonymous and nonsynonymous SNPs are summarized in Table 4 and Table S1-S6 in the supplemental material. An increase in detectable polymorphism was observed for phi6-E8G (paired t-test p= 0.001) after host shifting on to PA, but no significant change in SNP numbers was observed for phi6-WT and phi6-G515S (paired t-test p= 0.38, 0.40). The numbers of SNPs detected in the phi6-E8G population grown on PA were also significantly higher than that for the phi6-WT and phi6-G515S populations grown on PA (paired t-test p= 0.03, 0.0001 respectively). Gene p2 on the Large segment, coding for the RNA-dependent RNA polymerase, appears to maintain a constant, high level of diversity. The surprisingly large number of low-frequency SNPs for phi6-WT raised on PP demonstrated the potential of a dsRNA virus with a 13Kb length genome that grows ˜5 generations in overnight plaque growth to generate substantial genetic diversity. This may also be the reason phi6-WT is more able to readily infect PA with a high PA host range mutation frequency (Figure 5).

**Table 4.** Pairwise unique SNPs above 0.1% frequency in phi6 populations detected through deep sequencing using VarScan. S for synonymous, NS for non-synonymous change.

|  |  | WT |  |  |  | E8G |  |  |  | G515S |  |  |  |
| --- | --- | --- | --- | --- | --- | --- | --- | --- | --- | --- | --- | --- | --- |
|  |  | PP |  | PA |  | PT |  | PA |  | PT |  | PA |  |
|  |  | S | NS | S | NS | S | NS | S | NS | S | NS | S | NS |
| Non-Coding |  | 14 |  | 20 |  | 4 |  | 29 |  | 7 |  | 8 |  |
| Coding |  | 23 | 44 | 19 | 54 | 8 | 20 | 39 | 67 | 7 | 24 | 12 | 16 |
| Small | P8 | 2 | 4 | 0 | 1 | 0 | 1 | 2 | 3 | 0 | 0 | 2 | 0 |
|  | P12 | 0 | 1 | 8 | 14 | 0 | 1 | 4 | 5 | 1 | 0 | 1 | 4 |
|  | P9 | 0 | 3 | 3 | 7 | 0 | 1 | 4 | 5 | 0 | 0 | 1 | 0 |
|  | P5 | 2 | 4 | 0 | 1 | 0 | 1 | 2 | 3 | 1 | 1 | 0 | 2 |
| Medium | P10 | 0 | 1 | 0 | 2 | 0 | 0 | 1 | 1 | 1 | 0 | 0 | 1 |
|  | P6 | 0 | 2 | 1 | 1 | 1 | 0 | 1 | 6 | 1 | 3 | 1 | 1 |
|  | P3 | 7 | 13 | 5 | 17 | 0 | 2 | 11 | 21 | 0 | 3 | 6 | 6 |
|  | P13 | 0 | 1 | 0 | 1 | 0 | 0 | 1 | 1 | 0 | 0 | 0 | 0 |
| Large | P7 | 2 | 3 | 1 | 0 | 2 | 2 | 0 | 1 | 1 | 2 | 0 | 0 |
|  | P2 | 5 | 7 | 1 | 4 | 4 | 6 | 3 | 10 | 1 | 7 | 0 | 1 |
|  | P4 | 0 | 3 | 0 | 3 | 1 | 4 | 3 | 5 | 0 | 4 | 0 | 1 |
|  | P1 | 5 | 2 | 0 | 3 | 0 | 2 | 7 | 6 | 1 | 4 | 1 | 0 |

**Figure 5.**
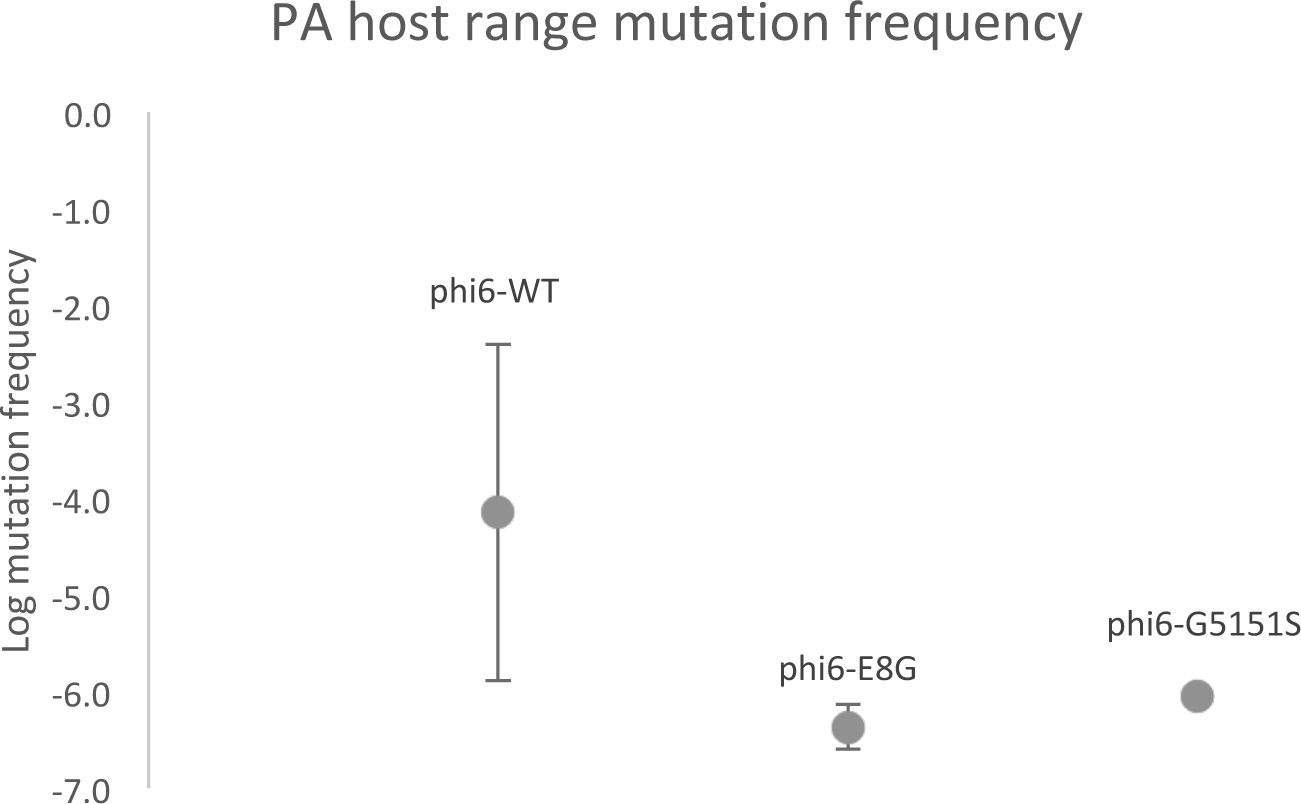
PA host range mutation frequency of phi6-WT, phi6-E8G, phi6-G515S. Values are average frequency of four purified single plaques, error bars are standard deviations.

### Relative fitness of naturally occurring phi6 PA host range mutants

A single mutation (A133V) was shared across the three genotypes, which prompted us to look at its fitness effects in the three genetic backgrounds. We used paired growth assays to measure the relative fitness of host range mutants on their shared, original host PP. Phi6-WT, the ancestor of all the tested strains, was the common competitor for all mutant genotypes and therefore has the relative fitness of 1 (Figure 6). Relative fitness was not affected when phi6-WT obtained A133V mutation on p3 (two-tailed one sample t-test, p= 0.66). However, when phi6-E8G and phi6-G515S gained the A133V mutation, each significantly increased their PP fitness on PP (Tukey’s HSD adjusted p= 0.000011, 0.011, respectively). K144R, on the other hand, was not beneficial in two genotypes on the PP host (p< 0.005).

**Figure 6.**
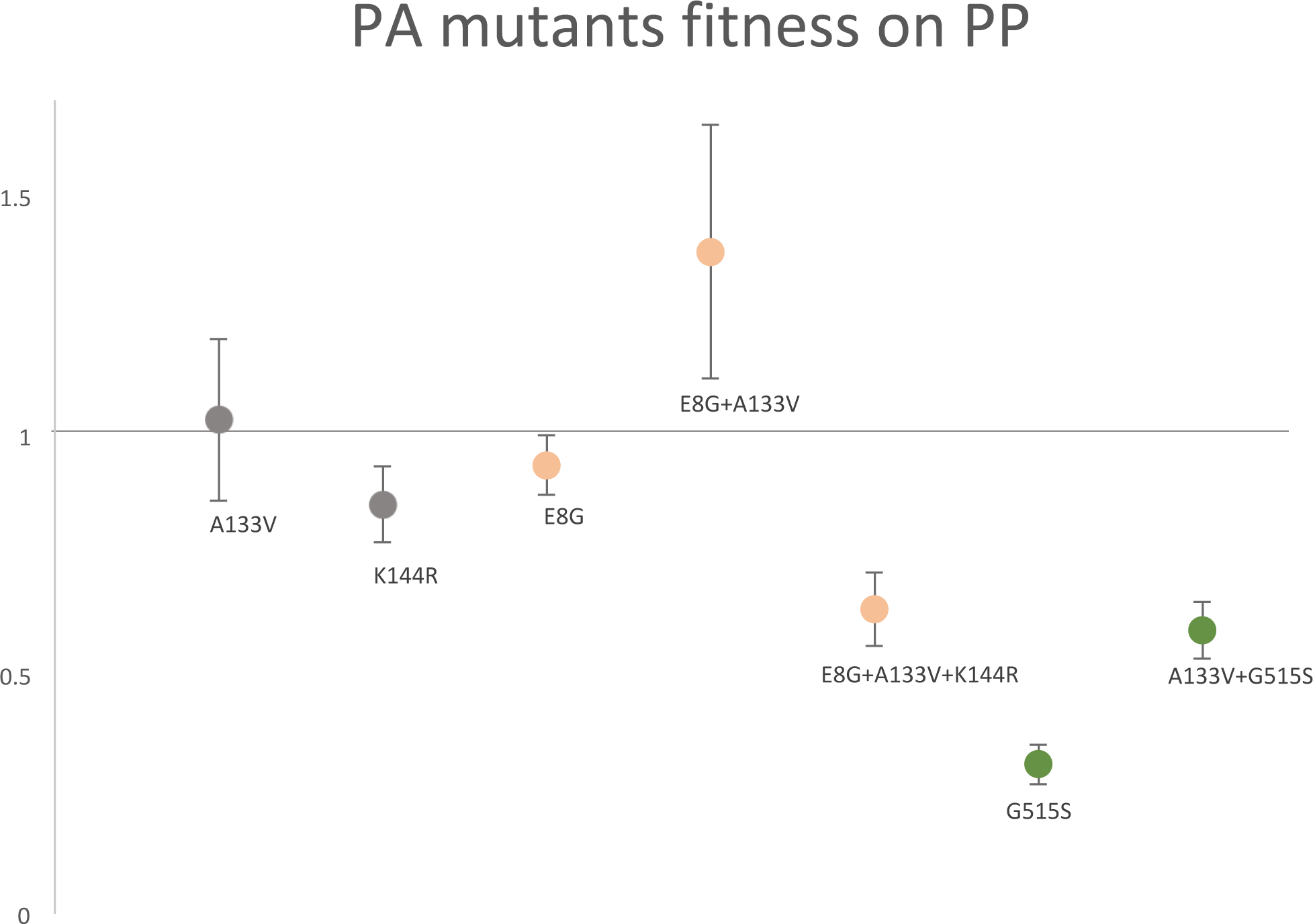
Relative fitness of host range mutants on PP. Same color genotypes share the same genetic background(phi6-WT: grey/black; phi6-E8G: peach; phi6-G515S: green). Values are average of six replicates, error bars are standard deviations.

## Discussion

We compared the host range mutational neighborhoods of wildtype phi6 to its isogenic host range mutants, and observed an epistatic constraint of existing host range mutations on the mutational neighborhood of subsequent novel host range expansion. Using next generation sequencing, we detected additional host attachment protein mutations missed in targeted Sanger sequencing of clones. We also identified – for the first time – secondary hot spots for host range expansion on the Small segment of phi6 genome.

Epistasis plays a large role in viral evolution, in part because viral proteins are often highly interactive, multifunctional, and many viral genomes are of limited size (36-38). The constraining effects of larger and smaller beneficial mutational neighborhoods were elegantly demonstrated in phi6 by Burch and Chao, who noted that the high mutation rate of phi6 was not sufficient to allow constrained genotypes to traverse a rugged fitness landscape (Burch and Chao, 2000). The ruggedness of viral fitness landscapes has been demonstrated for many viruses including HIV (37), Influenza virus A (39, 40), and Ebola virus (41), and the phenomenon of mutational neighborhoods constraining evolutionary trajectories has been demostrated in cellular organisms as well (42-44). Many of these studies involved prolonged experimental evolution, whereas our study used a very narrow window of lethal selection: a single night’s selection on a novel host. Nevertheless, we detected strong epistatic constraint from single amino acid changes, and in an ecologically realistic scenario for an RNA virus.

Epistatic interactions can exist between loci in the same gene, or at distant places in the genome (45, 46). We see both effects here – restricted PA mutations in P3, and elimination of the surprising Small segment mutations that can mediate PA host range in phi6-WT.

Evolvability may vary over evolutionary history (47), and we have only characterized one mutational step (PA mutation frequency) for these three phage genotypes. The genotypes have differences in mutational supply affected by either the size of the PA mutational neighborhood or population size (or both, (42)). Moderate differences in population size are a concern because phi6-WT was reared on the highly productive host PP and the two mutants were reared on PT, a host on which they are less productive. There are additional “maternal effects” on phi6 fitness due to host used to generate a high titer lysate, which could cause further differences in PA plaquing efficiency among phi6 strains grown on different hosts (23). However, the library preparation for Illumina sequencing involved the same amount of RNA, enforcing a similar population size of genomes sampled by sequencing. Phi6-WT had double the number of SNPs in high titer lysates in deep sequencing following double-plaque purification, demonstrating that constrained mutational neighborhood for the two host range mutant genotypes played a large role in the reduced evolvability on PA.

Further, our results do not necessarily mean that phi6-E8G and phi6-G515S are trapped on their fitness landscapes with regard to host range expansion on PA. If these mutants were allowed to evolve further (on the original host or within their novel host range), it is difficult to predict if their evolved descendants would face identical constraints when infecting the PA host – which more accurately reflects the serial host jumping we have observed for mammalian viruses (). It is easy to imagine that a phi6 P3 protein, somewhat destabilized by the addition of one host range mutation, cannot tolerate further destabilization while retaining its structure and function. The fitness benefits of A133V on the original host (PP) for both phi6-E8G and phi6-G515S may indicate that this is one of few (or the only) PA host range mutation in P3 that improves the mutants’ P3 structure and function. Other compensatory mutations acquired over evolutionary time could stabilize the P3 of descendants of phi6-E8G and phi6-G515S, creating larger mutational neighborhoods for PA host range expansion.

### Specific mutations found in our study

The most common means for a mutant to adapt is through additional (potentially compensatory) mutations rather than reversion of mutation (48). We observed three clones in the phi6-G515S population reverting (S515G) while fixing a PA host range mutation. This suggests that the G515S mutation is a relatively deleterious mutation to maintain in the genome, which is bolstered by the low fitness of phi6-G515S on PP (Figure 6).

While the Small segment has not previously been associated with host range, one of our PA host range mutations (P12:F176L) was previously observed in a phi6 evolution experiment on a different novel host (49). Changes in the Small segment-encoded protein P5 are known to affect phi6 thermal niche expansion (50), but this is the first time that membrane protein P9, enveloped lytic protein P5 and membrane morphogenic protein P12 (which is not found within the virion) were associated with host range. Across diverse viruses it is not uncommon for a variety of proteins in the envelope (such as P9) to interact with host receptor proteins (6, 51). It is more rare for non-structural genes that are not on the exterior of the virion to be a host range determinant, but there are examples of this: PB2 of avian influenza (52, 53) and in picornaviruses (54, 55). P5 plays a role in both phi6 entry and egress: copies of this muralytic enzyme underneath the lipid coat bore through the cell wall to allow phi6 to re-envelop itself with the host’s cytoplasmic membrane, and it is used to lyse host cells after sufficient replication (56-58). P12, however, is solely associated with egress from cells (forming membranes from the host cytoplasmic membrane around completed nucleocapsids) and is not detected in phi6 virions (35, 59). P12 was a hotspot of change in entropy for phi6-WT and 10 of the 13 clones that did not have a mutation in P3 had one of five nonsynonymous mutations in P12, which strongly suggests that this non-structural protein has a role in host range expansion to PA. Given our current understanding of P12’s role in the phi6 life cycle (34), this would require phi6-WT to be able to attach and infect PA but fail to show its infection through a plaque assay – and then mutations in a number of genes could boost the infectivity of phi6 to cause successful plaque formation. As phage host range is a difficult and debated phenotype to measure (60), and plaque formation is known to be affected by many genetic and environmental factors (61), this could well be the scenario for phi6-WT on hosts closely related to its original host PP, which express the same attachment site: the type IV pilus (62). It is known that DNA phage can attach to more hosts than they can productively infect, often due to successful host defense mechanisms such as CRISPR-Cas, restriction endonucleases and suicide of the infected cell prior to phage maturation (63); our RNA phage which are not known to trigger abortive infection or be susceptible to these defenses may still enter hosts they cannot productively exit. This suggests interesting follow up experiments with phi6-WT and hosts considered outside of its current host range. While spot plating is considered a sensitive method for detecting phage host range (64), lack of visible plaques does not definitively mean that phage cannot productively infect a given host at a low level or at a slow pace (65). It further suggests some of the host range mutation observed here may be better categorized as mutations that aid in spread among novel hosts rather than the attachment mutations that allow for a spillover infection, which are often considered separate steps in the emergence of a virus on a novel host (66). On a practical level, one or more of the PA-host range associated mutations in p12, p9 or p5 may prove useful as a Small segment marker for genetic crosses in phi6. The existing mutational markers on the Small segment are an easily reverted temperature sensitivity and an unstable genetic insertion (67, 68).

It remains unclear how the single PA mutant with no nonsynonymous mutation found in the genome (all genes Sanger sequenced) productively infects PA and forms plaques. Phi6 host range mutants without identifiable P3 mutations have been isolated in the past, but this is the first report of phi6 without any identified mutation from its immediate ancestor with a different host range. We can only speculate on the molecular processes underlying this plaquing – perhaps the match of lipid coats from those virions produced on PA have a higher infectivity on PA than those grown on PP (or other hosts, (23)).

In addition to phenotypic change without accompanying, explanatory genetic change, we also repeatedly observed some genetic change without an obvious cause or consequence. Sites at the single-stranded ends of phi6’s genomic segments experienced drastic changes in entropy, including at position 2843 of Small segment of phi6-WT and phi6-G515S, and position 2770 for all genotypes and position 3930 of Medium segment of phi6-WT and phi6-G515S. While the homologous 5’-ends are crucial for precise packaging, these positions are all at the 3’-end of the segments, which are important for RNA stability and polymerase recognition (69). It is unclear why these sites would become more or less diverse due to selection on PA.

### Contrasting clonal sequencing and population deep-sequencing

Next generation sequencing is now more frequently applied to microbial experimental evolution studies, changing how microbial populations are monitored and analyzed, focusing mostly on relative variant frequencies and their fitness effects (70-73). However, when determining population diversity structure, many studies still use cloning for isolate sequencing, and examining chromatograms to describe nucleotide polymorphism (74-76). Increasingly, studies have exclusively used deep-sequencing for population SNP detection (77, 78). In this study, both clonal sequencing and population deep sequencing had merits and shortcomings. Clonal mapping of mutational neighborhoods with 50 clones involved relatively small sample sizes but allowed unambiguous identification of single, double, and triple mutant combinations. Illumina sequencing of populations provided a more reassuringly complete picture of the mutational neighborhood – highly consistent with that of the clonal sequencing – but it might recover hitchhiking mutations that might not be responsible for the phenotype of interest, and cannot assign combinations of mutations to a single genome. For instance, more sophisticated approaches would be required to determine that the phi6 strains with Small segment mutations did not also have one or more of the prevalent p3 mutations, since these unconnected chromosomes could not become linked with longer sequencing reads. An additional durable problem is in haplotype determination. Population genetic analyses require a firm assessment of haplotypes in the population, and although many software programs are available for haplotype prediction, these programs were validated on shorter genomic regions and performed poorly on the tripartite 13kb phi6 genome, and accuracy depends heavily on read length, which we were unable to provide with 150bp Illumina single-end reads (79).

Using the change in Shannon entropy provided more accurate data analysis because it allowed us to cancel out a large amount of the noise produced by potential sequencing errors, and is ideal for our study’s purpose, which is contrasting populations before and after a challenge. However, we found we could not rely on entropy signals as estimates for mutations frequency or abundance. Although the top three most frequently observed P3 mutations in phi6-WT showed the largest change in entropy, this pattern does not apply to other observed host range mutations (e.g. phi6-WT P3: position 140, phi6-G515S P3: position 35, phi6-WT P9: position 8). Population deep sequencing provided an excellent snapshot of the mutational neighborhood, but it prevents many downstream analyses (including any further experimentation with clones of interest). However, it is cheaper and faster than clonal isolation and will serve the needs of many researchers, especially in studies of host range mutations for emerging disease surveillance.

## Acknowledgements

This research was funded by the National Science Foundation DEB 1453241 (to SD). We thank Nicole Wagner of the SEBS Genome Cooperative and Natasia Jacko for expert technical assistance. German Lagunas Robles and the students of South Brunswick High School’s 2016 Biotechnology class isolated several of the clones used in this study.

